# Assessing and characterising the repertoire of constitutive promoter elements in soil metagenomic libraries in *Escherichia coli*

**DOI:** 10.1101/211367

**Authors:** Cauã Antunes Westmann, Luana de Fátima Alves, Rafael Silva-Rocha, María-Eugenia Guazzaroni

## Abstract

Although functional metagenomics has been widely employed for the discovery of genes relevant to biotechnology and biomedicine, its potential for assessing the diversity of transcriptional regulatory elements of microbial communities has remained poorly explored. Here, we have developed a novel framework for prospecting, characterising and estimating the accessibility of promoter sequences in metagenomic libraries by combining a bi-directional reporter vector, high-throughput fluorescence assays and predictive computational methods. Using the expression profiling of fluorescent clones from two independent libraries from soil samples, we directly analysed the regulatory dynamics of novel promoter elements, addressing the relationship between the “metaconstitutome” of a bacterial community and its environmental context. Through the construction and screening of plasmid-based metagenomic libraries followed by *in silico* analyses, we were able to provide both (i) a consensus exogenous promoter elements recognizable by *Escherichia coli* and (ii) an estimation of the accessible promoter sequences in a metagenomic library, which was close to 1% of the whole set of available promoters. The results presented here should provide new directions for the exploration through functional metagenomics of novel regulatory sequences in bacteria, which could expand the Synthetic Biology toolbox for novel biotechnological and biomedical applications.

## INTRODUCTION

The study of prokaryotic transcriptional regulation is essential for understanding the molecular mechanisms underlying decision-making processes in microorganisms ^1^, comprising populational (e.g. colony structure, quorum sensing detection), ecological (e.g. nutrient acquisition, biomass degradation) and pathogenic behaviours (e.g. host recognition, biofilm formation). The activity of most bacterial promoters is usually dependent on the combined action of transcription factors and sigma factors in response to multiple environmental stimuli ^2^. For instance, in *Escherichia coli*, the compilation of decades of experimental data indicate that approximately 50% of its promoters are under the control of a single specific regulator, while all other genes are regulated by at least two transcription factors ^3^. Moreover, the recent development of experimental and large-scale sequencing techniques, together with powerful computational approaches have allowed both the discovery of insightful information about bacterial transcriptional systems and the development of novel approaches for studying those systems in higher depth ^4–7^. However, despite technical innovations, most of the studies are still centred on the model organism *E. coli*, a single bacterial species among at least 30,000 other already sequenced ^8^, in an estimated total of 1 trillion species ^9^.

With the advent of Metagenomics ^10^, the exploration of unculturable bacteria (approximately 99% of a bacterial community^11^ widely expanded genomic information, providing resourceful data about populational structures and genetic diversity in a myriad of environmental samples ^12–14^. Two main approaches are commonly adopted for those metagenomic studies ^15^: the sequence-based metagenomic approach, which relies on massive sequencing of metagenomic DNA and powerful bioinformatics tools for extracting information from the metagenomic sequences; and functional metagenomics ^16,17^, which directly explores the functionality of enzymes and other structural elements through a wide range of stress/substrate/product-based assays ^18–21^. In this context, although a large number of genes/ORFs has been discovered through the previously described approaches, the detection of novel bacterial regulatory elements using high-throughput technologies has been poorly explored, presenting so far a single well-defined method for the discovery of substrate-inducible regulatory sequences - SIGEX - ^19^ and a limited assay for exploration of constitutive promoters ^22^. This narrow range of methodologies is directly related to the biased functional search towards novel genes and to a lack of both experimental and computational tools for finding and validating promoter sequences in metagenomic libraries ^23^.

Unravelling novel bacterial promoters is essential for understanding the regulatory diversity of microorganisms, addressing important questions, such as the abundance of both constitutive and inducible elements in a metagenomic library, the bottlenecks regarding host choices (i.e. the constrains limiting the diversity of exogenous regulatory sequences that can be recognized by different hosts) and the correlation between promoter strength, transcriptional noise and the functional role of the regulated gene/operon ^23–26^. Furthermore, prospecting and characterising novel regulatory sequences is crucial for expanding the current Synthetic Biology toolbox and generating novel biotechnological applications. For instance, there is a high demand for novel constitutive and inducible promoters responding to process-specific parameters imposed by a wide variety of processes, such as industrial applications, heterologous protein expression and biosensors generation ^19,23,27–29^.

In this context, the most common strategy for prospecting regulatory sequences is the usage of unidirectional promoter trap-vectors, which consist in transcriptional fusions between DNA fragments and a reporter gene. This method has been widely employed for assessing regulatory sequences in genomic DNA ^30–33^, however its application in metagenomic DNA fragments has remained poorly explored^19^. The main constraint regarding the use of unidirectional systems is that bacterial genomes present a large variation in the percentage of their leading-strand genes, ranging from ~45% to ~90% ^34,35^. Thus, a bi-directional promoter reporter system would be preferable, by increasing the probability of finding promoter sequences. In the present work, we have developed a novel strategy for in-depth prospection, characterisation, and quantification of accessible promoter elements from soil metagenomic samples in *E. coli* as a standard host.

Although both constitutive and inducible promoters were potentially detectable by this method, we have focused exclusively on the study of the former, as a proof of concept, by avoiding substrate-based induction assays as previously reported ^18–21^. We have collected soil samples from two differentially biomass-enriched sites of a Secondary Atlantic Forest in South-eastern Brazil and generated metagenomic libraries in a bi-directional probe vector for primary screenings. We have characterised the expression behaviours of a large set of GFPlva expressing clones from both libraries and narrowed down our selection to 10 clones for an in-depth analysis regarding potential ORFs and endogenous promoters. By cross-validating *in silico* analyses and experimental data of predicted regulatory sequences, we have located and profiled the expression of 33 endogenous promoters within the selected clones (see Supplementary Table S2 online), providing resourceful information concerning the architecture and transcriptional dynamics of promoters from metagenomic fragments. Thus, in order to contribute to this set of accessible genetic features, we have used our gathered data to provide for the first time a direct estimation of the whole set of accessible constitutive promoters in a soil metagenomic library hosted in *E. coli*, which we have called the “metaconstitutome” of an environmental sample.

## RESULTS AND DISCUSSION

### Generating metagenomic libraries and screening for fluorescent clones

We have constructed and assessed two metagenomic libraries hosted in *E. coli* DH10B strain for the analysis of bacterial promoters in environmental samples (Figure 1). The libraries were generated from soil microbial communities of two sites bearing differential tree litter composition (*Anadenanthera spp.* and *Phytolacca dioica*) within a Secondary Atlantic Forest zone at the University of Sao Paulo, Ribeirão Preto, Brazil. Both metagenomic DNA were cloned into the pMR1 bi-directional reporter vector, which has GFPlva and mCherry reporter genes in opposite directions ^36^. Each metagenomic library presented about 250 Mb of environmental DNA distributed into approximately 60.000 clones harbouring insert fragments size ranging from 1.5 Kb to 7 Kb, with an average size of 4.1 Kb (Table 1). We have chosen fragments of 1.5-7 Kb in order to validate our strategy on standard-sized functional metagenomic libraries based on plasmid vectors ^18,19,37–39^. In total, 1,100 fluorescent clones, resulting in a rate of approximately one fluorescent clone every one hundred fifty clones (USP1) or every ninety clones screened (USP3), were manually selected under blue light exposition. Then, these fluorescent clones were directly recovered from LB agar plates supplemented with chloramphenicol. The direct screening was preferred over the use of metagenomic clone pools from stocks as it reduces the chances of both biased clone enrichment (e.g. clones with higher growth rates, usually clones bearing small inserts or without insert) and dilution of positive clones with impaired growth (e.g. clones with high expression of GFP and/or other exogenous genes), avoiding thus clonal amplification.

**Table 1.** Features of the generated metagenomic libraries.

| Metagenomic Library | USP 1 | USP 3 |
| --- | --- | --- |
| Total number of clones | 100,000 | 90,000 |
| Percentage of clones with insert | 60% | 70% |
| Number of clones with insert | 60,000 | 63,000 |
| Total number and rate* of fluorescent clones | 400 (1:150) | 700 (1:90) |
| Total number and rate* of green clones | 270 (1:220) | 400 (1:157) |
| Total number and rate* of red clones | 130 (1:460) | 300 (1:210) |
| Average insert size | 4,5 kb | 3,7 kb |
| Library Size | 270 Mb | 233 Mb |
| Estimated number of genomes** | 60 | 52 |

\* Rate represented by the number of fluorescent clones divided by the total number of clones with inserts.
\*\* Assuming 4.5 Mb per genome <sup>81</sup>.

**Figure 1.**
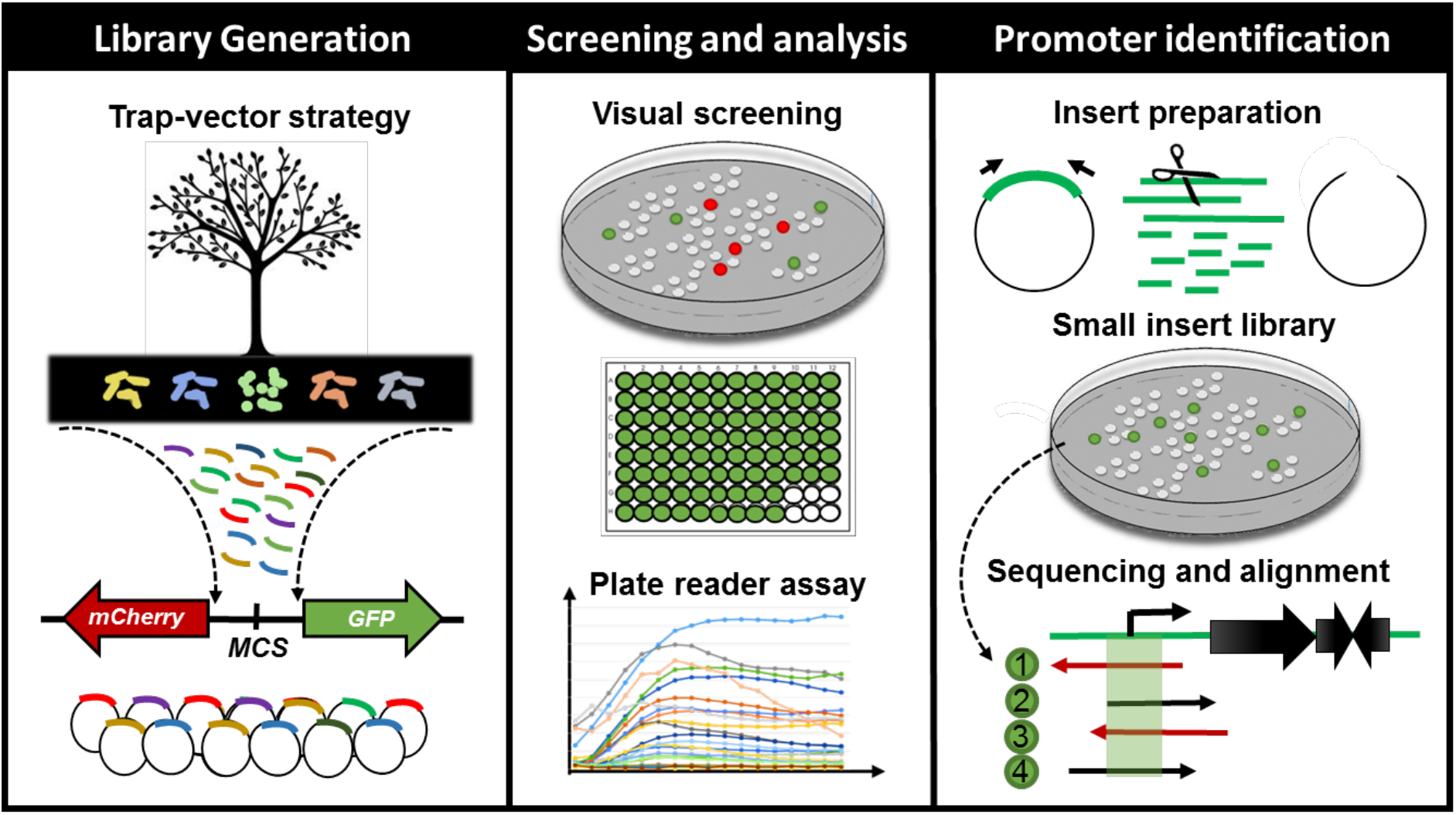
Schematic representation of the workflow for finding, characterising and cross-validating novel bacterial cis-regulatory elements in environmental samples. From left to right: firstly, we have generated metagenomic libraries from soil samples in *E. coli* DH10B. The DNA fragments were cloned into a bi-directional reporter trap-vector (bearing *mCherry* and *GFPlva* fluorescent reporters), pMR1, which allowed for the screening of promoters in both DNA strands. Secondly, we have manually screened all visible fluorescent clones from our metagenomic libraries and analysed the expression patterns of all green fluorescent clones on a microplate reader during 8 hours. Lastly, we have selected ten clones based on their GFPlva expression patterns for an in-depth analysis combining experimental (small DNA insert library generation) and *in silico* promoter prediction. This integrated strategy has allowed us to identify, validate and estimate the accessibility of novel promoter regions from metagenomic libraries.

### Evaluating the expression dynamics of fluorescent clones

In order to analyse the expression patterns of the isolated clones, we evaluated the intrinsic dynamics of GFPlva and mCherry by randomly selecting 20 clones expressing each reporter (as schematically represented in Figure 1). As represented in Figures 2A-B, we found that clones expressing mCherry were not suitable for standard microplate 8 hour assays, as the fluorescence intensity values differed dramatically between 8 and 24 hours after the beginning of the experiment. The slow kinetics of mCherry expression has already been reported as a consequence of a two-step oxidation process for protein maturation when compared to the one-step maturation process found in GFP reporters ^40^. On the other hand, the clones expressing GFPlva presented the enhanced intrinsic properties for microplate assays, supported by the observation of very similar fluorescence intensities between the two time points tested. Furthermore, the GFPlva has an LVA-degradation tag attached to its C-terminal, which reduces GFP accumulation and increases protein turnover, generating a more precise fluorescence output on analysis of expression patterns ^41^.

**Figure 2.**
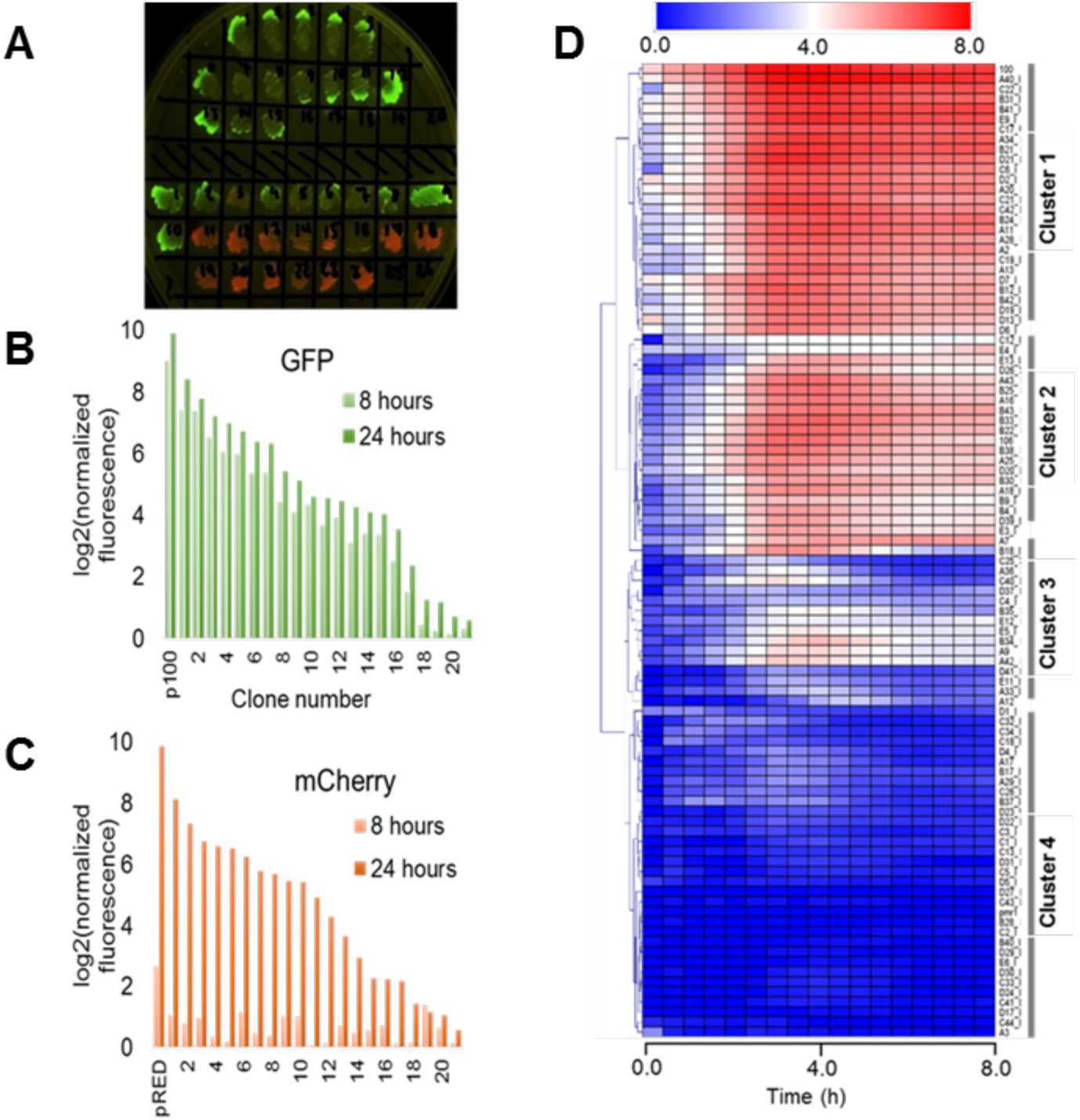
Evaluating the expression dynamics of fluorescent clones. **A)** LB-agar plate under blue light excitation comprising a subset of metagenomic isolated clones expressing GFPlva (top) and mCherry (bottom) fluorescent reporters. A few clones were observed to express both reporters. All isolated clones were initially considered to hold at least one endogenous promoter. **B-C)** Indirect assessment of maturation times from both fluorescent reporters GFPlva (B) and mCherry (C) after 8 hours (light bars) and 24 hours (dark bars) of the beginning of the experiment. Maturation times are substantially lower for mCherry than for GFPlva, which excluded the former from further analyses. Positive controls for GFP and mCherry are represented by p100 and pRED, respectively. Fluorescence data has been normalised by OD_600_ values for each sample following normalisation by values from the negative control (empty-pMR1). Data was transformed to log2 scale to allow better visualisation of fluorescence variation. **D)** Hierarchical representation of a metaconstitutome (i.e. all expression profiles from a single metagenomic library. Fluorescence time-lapse dynamics were measured during 8 hours for each clone and represented as heat maps. Promoter activities (calculated as GFP/OD600) were normalised by the negative control (*E. coli* DH10B harbouring empty pMR1) and transformed to log2 scale in order to facilitate the visualisation of subtle activities. Data are representative of three independent experiments.

Thus, 260 clones expressing GFPlva (160 clones from the USP1 library and 100 from USP3) were selected for further analysis of expression patterns on microplate reader assays with biological and technical triplicates. The dynamic profiles for each clone were converted into heat maps and hierarchically clustered by a Euclidean Distance algorithm into a dendrogram, concisely representing the expression patterns of each metagenomic library. In order to assess the diversity of promoter strengths among the generated metagenomics libraries, three previously characterized constitutive promoters (see Experimental Procedures for further information) positioned upstream a GFPlva reporter were used as standards for strong, medium and weak expression profiles (referred here as p100, p106 and p114, respectively). Considering both metagenomics libraries, we have found a total of 30 strong promoters showing a strength similar to the p100 control, 40 medium strength promoters similar to the p106 control, 60 weak promoters similar to the p114 control and a wide range of promoters with particular expression patterns which did not cluster with any of the previously mentioned positive controls (Figure 2C and Supplementary Fig. S1 online). Since the exploration of distinct expression behaviours is essential for expanding the current set of commercial promoters, the diversity of expression profiles highlighted in this study has supported the current framework as a promising strategy for finding novel promoters for downstream applications.

Furthermore, concerning the hierarchical organization of the expression profiles, the dendrogram of the USP3 library (Figure 2C) suggests the presence of at least four well-defined expression clusters comprising: (i) high, (ii) medium, (iii) low and (iv) very low expression profiles. A very similar pattern was identified in the expression dendrogram independently generated for the USP1 metagenomic library (see Supplementary Fig. S1 online), suggesting those clusters might be depicting broader trends of organizational expression patterns in nature. Independent studies on microbial communities from aquatic environments have described similar patterns by evaluating gene expression through metatranscriptomic analysis ^42–45^, indicating that our observations are not restricted to the assessed soil samples. However, further studies with a systematic application of the methodologies described here over a broader range of environmental samples would be required for evaluating these profiles.

### In silico analysis of DNA metagenomic fragments from selected clones

From the 260 assessed samples, we have selected 10 clones displaying particular profiles (see Supplementary Fig. S2 online) depicting the diversity of expression behaviours found in both libraries. The inserts from selected clones were sequenced and analysed for both potential ORFs and RpoD-related promoter regions (−10 and −35 conserved regions). In the case of the identification of putative genes, twenty-nine ORFs with significant *E-values* (<0,001) were found (Table 2 and Supplementary Table S1 online) unevenly distributed between both DNA strands, in line with a lack of strong directional trends regarding bacterial genome organization ^46^. The ORFs were also classified within a range of functional classes (delineated by MultiFun^47^ and potential bacterial phyla (see Supplementary Fig. S3 online). For this, we carried out the analysis of the microorganisms associated with the closest similar protein of the identified ORFs (Table 2). The most abundant ORFs were related to unknown functions (31%) and metabolism (31%), followed by stress adaptation cell processes (17%) (see Supplementary Table S1 online), while the most abundant phyla related to the recovered ORFs were Proteobacteria (35%), followed by Bacteroidetes (22%) and Chloroflexi (14%) (see Supplementary Fig. S3 online). The relative abundance of the guanine-cytosine content of each insert was also assessed (Table 2), resulting in a median of 54%, varying from 43% to 61%, indicating their diverse phylogenetic affiliation. These results are in agreement with previous G-C content diversity analyses of soil samples which ranged from 50% to 61% ^48–50^. Even with a limited sample size when compared to NGS-based metagenomic studies, the abundance of gene functions and bacterial groups predicted in this work was similar to the ones found in previous studies in soil microbial communities ^51–53^. Considering the above, these results suggest that different bacterial groups could be the sources of accessible promoters in *E. coli*, that is, regulatory sequences recognizable by the molecular transcriptional machinery of *E. coli* that allowed the expression of the reporter genes.

**Table 2.**
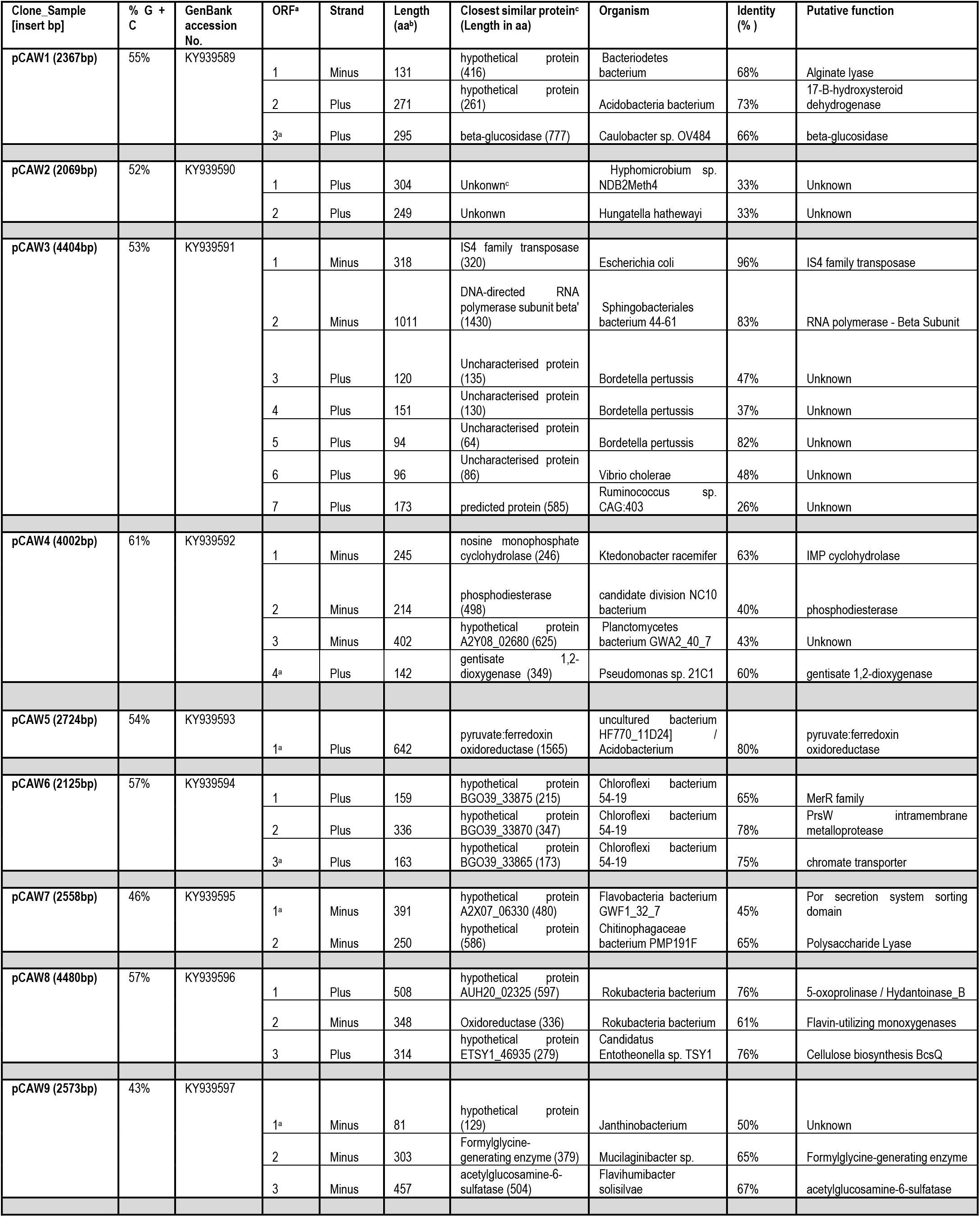
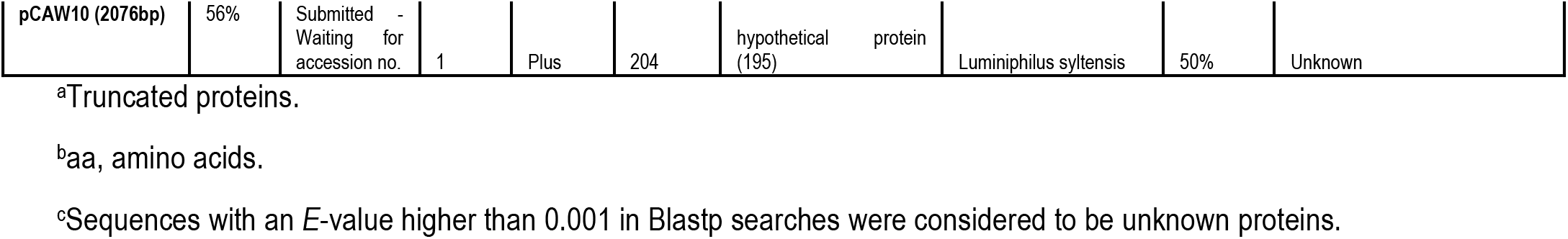
Description of the ORFs contained in plasmids from the selected clones (pCAW1 to pCAW10) and their sequence similarities.

The *in silico* promoter prediction has also provided relevant information concerning the potential number of regulatory regions on each selected fragment. The BPROM software ^54^ has been extensively employed in other promoter prediction studies and is based on the analysis of the −35 and −10 consensus sequence of RpoD promoters. The main sigma subunit, sigma-70 encoded by *rpoD*, plays a major role in transcription of growth-related genes, the so-called housekeeping genes ^55–57^. From the *in silico* analysis, a total of 140 promoters were predicted among the 10 selected clones, suggesting an average of 5 RpoD-related promoters/Kb. This led us reasoning that most of the expression profiles previously described (Figure 2C and Supplementary Figure S1 online) were representing the dynamics of the merged promoters present in the metagenomic fragment. Considering that, we delineate a strategy to experimentally assess the number and location of accessible promoters from our selected clones, contrasting experimental results with *in silico* data.

### Experimental identification, characterisation, and cross-validation of promoter regions

In order to explore the potential set of accessible promoter regions from our metagenomic libraries, we developed a small DNA insert library generation approach (Figure 1). Firstly, the plasmids from the previously 10 selected clones (original clones) were pooled together for insert amplification in a single PCR reaction. The resulting amplicons were fragmented by Sau3AI digestion and DNA fragments ranging from 0.2 Kb to 0.5 Kb were selected for subsequent cloning into the pMR1 vector. The generation of this sub-fragment library allowed the screening for both red and green fluorescent colonies as they would represent the accessible set of promoters among the metagenomic DNA fragments studied. It is important to highlight that as the cloning process was not directed, small fragments bearing promoter regions had a 50% chance of getting cloned in any direction, thus clones expressing mCherry were also isolated for subsequent sequencing. A total of 100 clones coming from the small DNA insert library (80 expressing GFPlva and 20 expressing mCherry) were sequenced and then align against the original metagenomic fragments. As a result, we have identified at least 33 promoter regions within the initial set of the selected metagenomic clones (Figure 3, Supplementary Fig. S4 and Supplementary Table S2 online). These findings showed that the *in silico* prediction of 140 RpoD-related promoters was overestimated in comparison with the experimental results. The above can be explained since prediction algorithms usually misrepresent nature by underestimating or overestimating results due to a lack of information regarding diversity and variability of natural *cis*-regulatory sequences ^58–60^.

**Figure 3.**
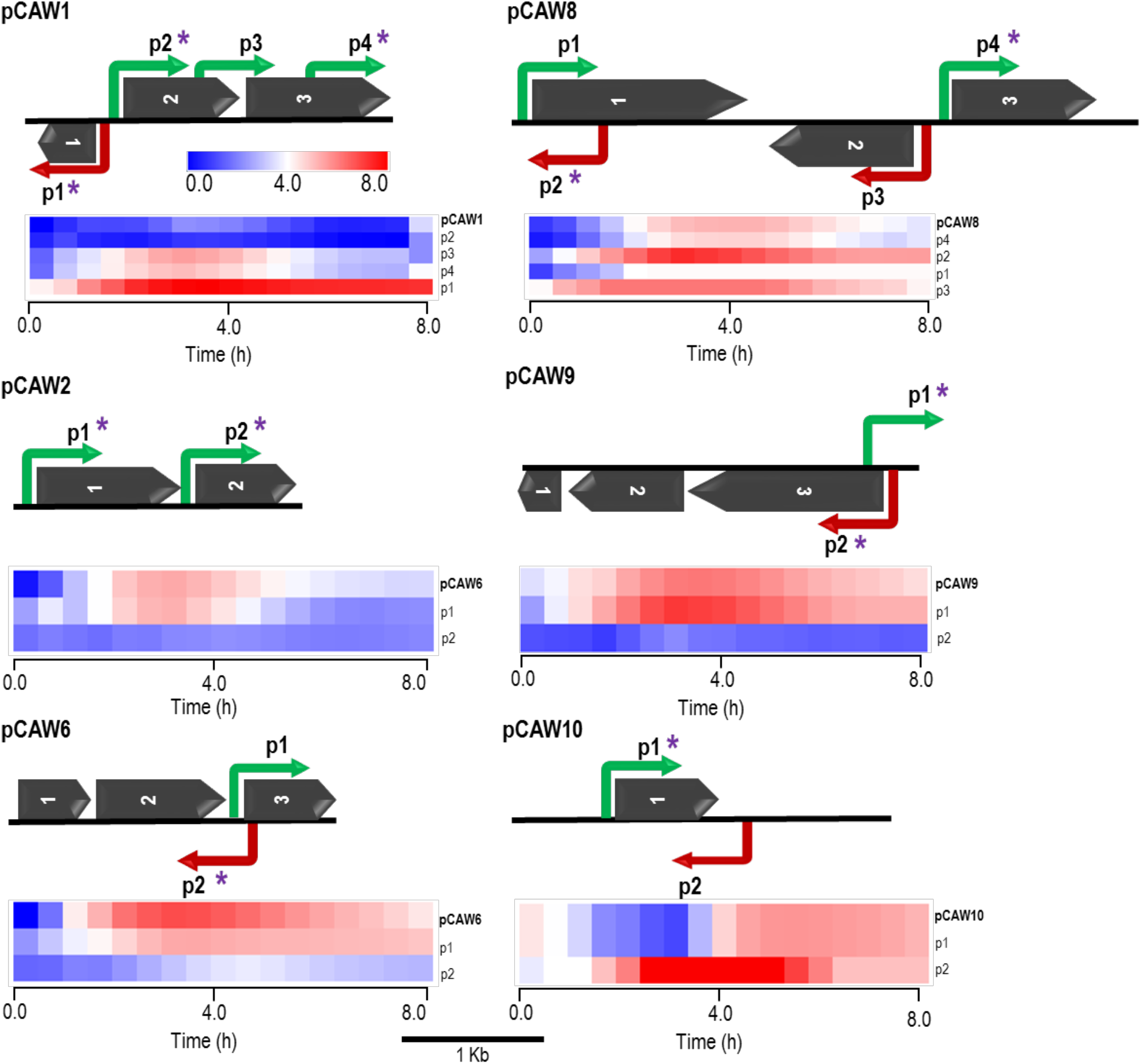
Schematic representation of six metagenomic inserts (contigs) showing predicted ORFs and experimentally validated/characterised promoters. Each contig is identified on the far left of each subfigure. Promoters are indicated by elbow-shaped arrows and name according to their relative position in the contig. Promoter directionality, regarding the leading and lagging strands, is represented by green and red colours, respectively. Asterisks over specific promoters indicate regulatory regions which were cross-validated by matching *in silico* predictions. Dark arrows represent predicted ORFs, according to their relative positions in each contig (see Table 2 for more information). All genetic features respect their original relative sizes, following the 1 Kb scale depicted at the bottom of this figure. Beneath each metagenomic insert, there is a heat map cluster representing the whole set of promoter activities measured during 8-hours fluorescence assays. The first line of each cluster shows the original expression profile initially measured for each metagenomic insert. All other lines represent expression activities from de novo experimentally validated promoters within each contig (small DNA fragments). The second line of each cluster represents the endogenous promoter showing the most similar activity with respect to the original expression profile for each contig. All expression profiles are properly identified at the most rightmost side of each line, following their respective contig/promoter name. For the supplementary set of analysed contigs, see Supplementary Figure S4 online.

Additionally, the current experimental approach allowed us not only to identify novel promoter regions but also to determine promoter directionality. The evaluation of promoter localization within the 10 selected clones revealed that from the 33 experimentally selected small fragments, 7 (21%) were considered intragenic promoters while the remaining 79% (26 promoters) were considered primary promoters, defined as the furthest upstream promoter in a gene/operon ^61^. This small-scale analysis slightly diverges from architectural features found in *E. coli* K-12 genome in which the promoter dataset was dominated by primary promoters (66.3%), with a lower number of secondary promoters (19.6%), defined as intergenic and downstream of primary promoters ^61^, internal promoters that are intragenic (9.8%), and antisense (4.2%) promoters ^61,62^. This observation might reflect the diversity of genomic architectures in metagenomic libraries and highlight the current underestimation of bacterial intragenic promoters, which doubled the number in comparison to *E. coli*.

Based on the alignment results, we selected a defined set of small fragment clones related to each original sequence for dynamic expression profiling on a microplate reader. The results showed that for each set of small-fragments belonging to a DNA metagenomic clone, there was at least one with an expression pattern corresponding to the original clone previously observed (Figure 3 and Supplementary Fig. S4). Similarly, we identified other clones bearing small-inserts with individual profiles different to the primarily observed, representing alternative promoter regions in the original sequence that were not mapped in the initial approach (Figure 3). The diversity of the promoter expression profiles found in a single original metagenomic clone has a multifactorial nature, ruled by different processes. Firstly, it should be considered the inherent relationship between the regulatory dynamics and the functional role of the regulated gene ^26^. Secondly, the transcriptional bias imposed by the *E. coli* molecular machinery, which would recognize orthologous sequences, but not necessarily reproduce the original behaviours found in natural hosts ^23,39,63,64^. Finally, another point to be considered is that the increase in expression levels can be the result of the artificial juxtaposition of the promoter to the fluorescent reporter ribosome binding site, as a consequence of the cloning process.

Regarding *in silico* cross-validation, from the 33 experimentally validated promoters, 23 RpoD-related promoters (70%) were supported by the algorithmic analysis as they were aligned to their respective original sequences (Figure 3). On the other hand, the remaining 10 sequences (30%) were considered as promoters exclusively identified by experimental approaches. We hypothesized that these sequences could be either recognized by other sigma factors than sigma70 or presented unusual consensus sequences for −10 and −35 boxes which has bypassed the algorithmic analysis. However, experimental validation in *E. coli* strains lacking diverse sigma factors genes should be necessary for a more accurate conclusion.

Finally, sequences of the above experimentally validated promoters were characterised accordingly to previous studies reported in the literature. For this, we adopted an *in silico* classification proposed by Shimada et al ^65^ (2014), in which constitutive promoters present a high-level conservation of the consensus sequence for the major sigma factor RpoD, that is, the elements TTGACA (-35) and TATAAT (-10) separated by approximately 17 bp (Figure 4A and B). Constitutive promoters are defined as promoters active *in vivo* in all circumstances, and, on the other hand, inducible promoters are switched ON and OFF by transcription factors depending on the *in vivo* conditions ^65^. The Logo pattern ^66^ generated from the alignment of the 33 identified metagenomic promoters (Figure 4C) indicated that positions −35 and −34 (−35 box) and positions −8, −7 and −3 (−10 box) were highly conserved. Although this logo pattern was distant from the proposed for the RpoD-dependent constitutive promoters identified *in vitro* (Figure 4A ^65^), was very similar to previously described consensus ^67^ from experimentally validated promoter sets from RegulonDB ^3^ and EcoCyc ^68^ databases (Figure 4B). To conclude, the results presented here has allowed us to identify a consensus for exogenous promoter recognition in *E. coli*, which can be an important resource for defining host-dependent restrictions in functional metagenomics.

**Figure 4.**
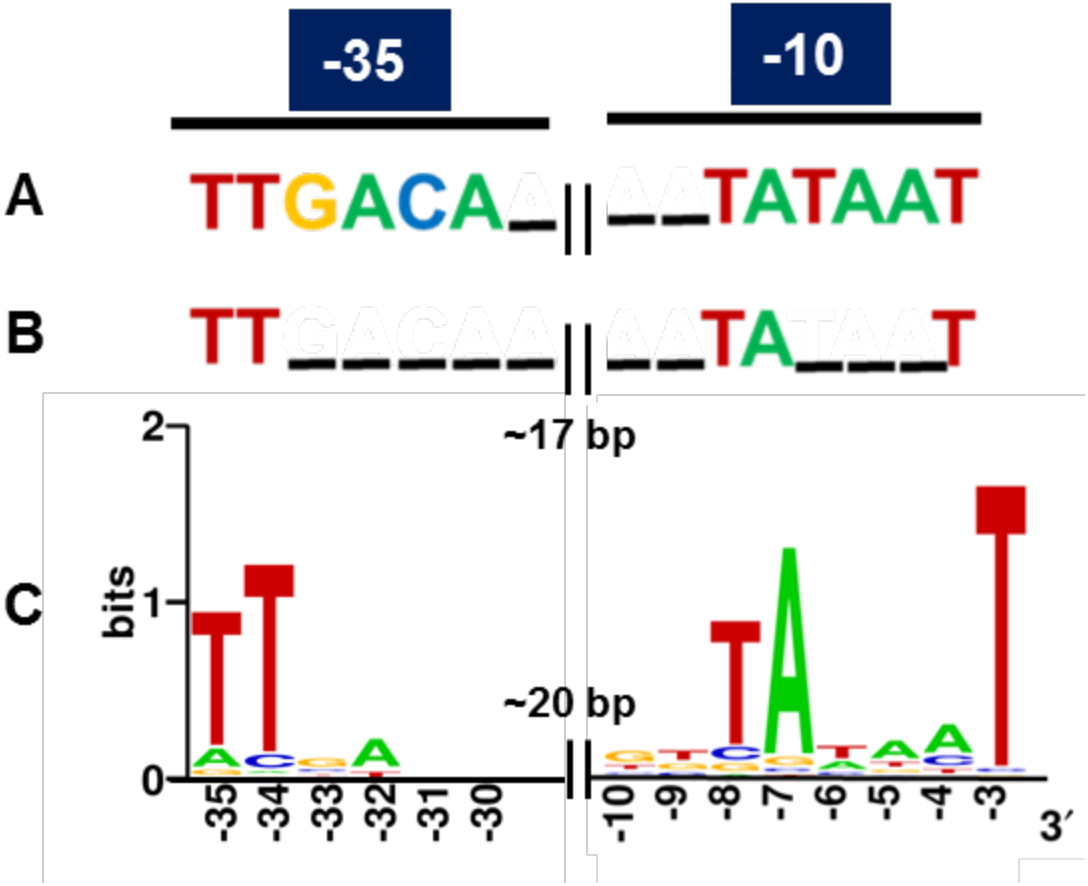
Consensus of RpoD-related metagenomic promoters. **A)** Known consensus sequences of the RpoD-dependent promoter determined in vitro, TTGAAC (-35) and TATAAT (-10) separated by 17 plus/minus 2 bp in E. coli ^65^. **B)** Known consensus sequences of 582 promoters experimentally validated in E. coli ^3,65,68^. **C)** The sequences of the 33 promoters experimentally validated in this study were aligned and subjected to Logo analysis ^66^. The consensus from the metagenomic set (C) is very similar to the one from the experimentally validated set from E. coli (B).

### Estimating the accessibility of promoters in random metagenomic libraries

In the present work we provide, for the first time in literature, a quantitative estimation regarding the accessibility of natural promoters in random metagenomic libraries, supported by the integration of both *in silico* and experimental results. We have estimated the existence of at least 553,300 promoters virtually recognized by *E. coli* in a standard functional soil metagenomic library, from which approximately 4,961 promoters (~1%) were readily accessible by our methodologies (see Experimental Procedures for calculations details). For the sake of comparison, we have estimated an average rate of 1.1 promoters/Kb potentially recognizable by *E. coli* host in our metagenomic fragments, which seems reasonable when compared to the rate of reported promoters in the well-studied genome of *E. coli* K-12, ranging from 0.5 to 2.7 promoters/Kb, depending on both chosen datasets and *in silico* prediction parameters ^61,62,65,69^. We have also assessed the genomic features from 24 bacterial species catalogued on the DOOR2 database (Database of prOkaryotic OpeRons) ^70^ for a broader promoter rate estimation, resulting in 2 promoters/Kb, which is also in concordance to our estimation and to others reported in literature ^61,62,65,69^. It should be mentioned that the average promoter rate from the present study (1.1 promoters/Kb) is probably an underestimation of the whole set available as it is restricted to clones expressing GFP under specific experimental conditions. Consequently, modifications of laboratory conditions during the screenings (such as growth-phase, temperature, exposure to different chemicals and substrates, to cite some) would probably reveal novel promoter elements ^71^.

A seminal study in functional metagenomics provided by Gabor *et al* ^63^(2004) estimated on a theoretical basis, using 32 prokaryotic genomes, that 40% of the enzymatic activities present in a soil metagenomic library could be readily accessed using *E. coli* as a host in an independent gene expression mode (in which both the promoter and the ribosome binding sites (RBS) are provided by the metagenomics insert). Moreover, it was predicted that Firmicutes, instead of Proteobacteria, would present the largest fraction of independently expressible genes (73%). Contrastingly, recent empirical studies on *E. coli* and other hosts have shown that functional expression faces a myriad of challenges that were not taken into account in previous mathematical models, such as codon usage, improper promoter and RBS recognition ^72^, missing initiation factors, protein misfolding, missing co-factors, breakdown of product; improper secretion of product, toxicity of product or intermediates and formation of inclusion bodies ^24,25^. Since it is impossible to predict the effect of the previously described difficulties in unknown metagenomic fragments, the actual fraction of genes that can be successfully expressed in *E. coli* is probably significantly lower than the proposed by Gabor and collaborators^63^ (2004). In this context, our work supports the previous arguments ^24,25^ highlighting the large gap between theoretical predictions and experimental data as we have shown only a small portion of the whole set of promoters is accessible for *E. coli* in metagenomics libraries (~1%). Thus, we stress the importance of feeding mathematical models with empirical data in a continuous iterative process for improving its predictive power.

## CONCLUSIONS

In summary, we have developed a novel methodology for prospecting, characterising and estimating the accessibility of promoter sequences in metagenomic samples by combining experimental and *in silico* approaches. The expression profiling of fluorescent clones was used for the first time as a direct approach to analyse the regulatory dynamics of an environmental sample, bearing great potential for revealing insightful trends regarding the transcriptional diversity of microbial communities. It has already been computationally demonstrated by Fernandez *et al.* (2014) ^73^ that the microbial metaregulome – the whole set of regulons of an environmental sample – is shaped by the physicochemical conditions of the environment as an adaptive process. Thus, future studies systematically applying our methodology to a range of environmental samples will greatly contribute to understanding this relationship between regulatory diversity and environmental adaptation in bacteria. At the same time, it can also be further applied to the design of efficient microbial communities for therapeutic or ecological needs ^73–76^.

Through the generation of a small-DNA insert library approach combined to *in silico* promoter prediction we were able to provide both (i) a consensus of recognizable exogenous regulatory sequences in an *E. coli* host and (ii) an estimation of the accessible promoter sequences in a plasmid-based functional metagenomic library, which was close to 1% of the whole set of available promoters. These are resourceful data for building a concise framework regarding the accessibility of genetic features from metagenomic libraries and how it can be influenced by the choice of different microbial hosts ^23,63,64^ or by the tinkering of the host’s transcription systems ^72,77,78^.

Although this work provided seminal information regarding promoter accessibility in metagenomics libraries, further high-throughput studies optimizing the proposed methods (e.g. application of automated screening methods; exploration of the whole set of fluorescent clones in a metagenomics library by Next-Generation-Sequencing) will be essential for expanding our current estimation into a more holistic landscape. Finally, we highlight that besides providing novel approaches for studying the regulatory diversity underlying environmental microbial communities, this work should be extremely useful for expanding the current Synthetic Biology toolbox through the discovery and characterisation of novel regulatory features.

## EXPERIMENTAL PROCEDURES

### Bacterial strains, primers, plasmids and general growth conditions

*E. coli* DH10B (Invitrogen) cells were used for cloning and experimental procedures. *E. coli* strains were routinely grown at 37ºC in Luria-Broth medium or M9 minimal medium ^79^ (6.4 g/L Na2HPO4·7H2O, 1.5 g/L KH2PO4, 0.25 g/L NaCl, and 0.5 g/L NH4Cl) supplemented with 2 mM MgSO4, 0.1 mM casamino acid, and 1% glycerol as the sole carbon source. When required, chloramphenicol (Cm) (34 µg/mL) was added to the medium to ensure plasmid retention. When cells were grown in minimal medium, antibiotics were used at half concentrations. Transformed bacteria were recovered on LB (Luria–Bertani) liquid medium for 1 hour at 37°C and 180 r.p.m, followed by plating on LB-agar plates at 37°C for at least 18 hours. All constructions were cloned into the pMR1 bi-directional-reporter vector ^36^, which carries mCherry and GFPlva, a short-lived variant of GFP.

### Metagenomic libraries construction and screening for fluorescent clones

The metagenomic libraries used in this work were generated in our laboratory from two soil samples of a Secondary Atlantic Forest at the University of Sao Paulo, Ribeirão Preto, Brazil. Each sample was differentially enriched regarding tree species abundance on plant-litter composition: (i) enriched in leaves from *Phytolacca dioica* and (ii) from *Anadenanthera spp.* DNA was extracted from soil samples using the UltraClean™ Soil DNA isolation Kit (Mo Bio Laboratories, Solana Beach, CA, USA). For the construction of the libraries, metagenomic DNA was partially digested using Sau3AI, and fragments from 1.5 kb to 7 kb were extracted from an agarose gel for ligation into the dephosphorylated and BamHI-digested pMR1 vector. Ligation mixtures were transformed by electroporation into *E. coli* DH10B cells. To amplify the libraries, they were grown on LB agar plates containing Cm and incubated for 18 h at 37°C. Both green and red clones were manually isolated from LB-agar plates exposed to blue light wavelength (at approximately 470 nm) by a transilluminator (Safe Imager™ 2.0 Blue Light Transilluminator). Ten fluorescent and twenty non-fluorescent clones were randomly picked from each library and had their plasmids extracted, following digestion with EcoRI and SmaI enzymes for checking presence/absence of inserts and their sizes. Cells from the same library were collected and pooled together in LB supplemented with 10% (wt/vol) glycerol for storing at −80°C. The plasmids from the 10 selected clones were isolated from individual clones and transformed into new *E. coli* DH10B cells to reconfirm expression patterns.

### Nucleic acid techniques

DNA preparation, digestion with restriction enzymes, analysis by agarose gel electrophoresis, isolation of DNA fragments, ligations, and transformations were done by standard procedures (Ausubel et al., 1994). Plasmid DNA was sequenced on both strands by primer walking using the ABI PRISM Dye Terminator Cycle Sequencing Ready Reaction kit (PerkinElmer) and an ABI PRISM 377 sequencer (Perkin-Elmer) according to the manufacturer’s instructions.

### GFP fluorescence assay and data processing

To measure promoter activity, freshly plated single colonies were grown overnight in M9 medium supplemented with required antibiotics. Samples were diluted 1:20 (v/v) in M9 medium for a final volume of 200uL in 96-well microplates. Cell growth and GFP fluorescence were quantified using a Victor X3 plate reader (PerkinElmer, Waltham, MA, USA). Promoter activities were expressed as the emission of fluorescence at 535 nm upon excitation with 485 nm light and then normalised with the optical density at each point (reported as fluorescence/OD600) after background correction. Background signal was evaluated with non-inoculated M9 medium and used as a blank for adjusting the baseline of measurements. *E. coli* DH10B harbouring the pMR1 empty plasmid was used as a negative control. Three different positive controls were used, consisting in *E. coli* DH10B harbouring pMR1 plasmid with one of the following synthetic constitutive promoters from the iGEM BBa_J23104 Anderson’s catalogue (http://parts.igem.org/Promoters/Catalog/Anderson) ^80^ upstream a GFPlva reporter: J23100, J23106 and J23114 (referred here as p100, p106 and p114, respectively). Unless otherwise indicated, measurements were taken at 30 min intervals over 8 h. All experiments were performed with both technical and biological replicates, being biological triplicates evaluated as independent measurements on different dates. Raw data were processed and plots were constructed using Microsoft Excel. All data was normalised by background values and transformed to a log2 scale for better data visualisation. Heatmap dendrograms with expression profiles were generated by using MeV2 (http://mev.tm4.org/) software.

### Small-DNA inserts libraries generation and screening

In order to experimentally find and validate the promoter regions from each of the ten selected metagenomic clones, an experimental technique was developed based on the previously described methodology of metagenomic library construction. All selected clones had their plasmids extracted and pooled together in an equimolar ratio. The pooled sample was amplified through a single PCR reaction using high-fidelity polymerase enzyme (Phusion) and previously described primers flanking the MCS region (Multiple Cloning Site) of the pMR1 vector, into which the metagenomic inserts were cloned. The resulting amplicons were firstly submitted to an analytical digestion followed by electrophoretic analysis for finding the optimal concentration of Sau3AI enzyme for obtaining fragments size ranging from 0.1Kb to 0.5Kb. Then, the purified pooled samples were fragmented by Sau3AI in preparative digestion and thereafter punctured from a 1% agarose gel in the region between 0.1 Kb and 0.5 Kb. These small DNA fragments, in turn, were ligated to pMR1 vector. Aliquots of electrocompetent *E. coli* DH10B cells were transformed with ligated DNA. A total of 100 fluorescent clones (80 expressing GFP and 20 expressing mCherry) were isolated under blue light excitation screening and had their plasmids extracted for sequencing reactions. Fluorescent clones were stored at −80°C in LB medium supplemented with required antibiotics and 10% glycerol (v/v).

### In silico analysis of ORFs and promoter regions

The inserts of selected clones were sequenced on both strands as previously described. Sequences were manually assembled for the generation of 10 contigs. Putative ORFs were identified and analysed using the online ORF Finder platform, available at the NCBI website (http://www.ncbi.nlm.nih.gov/gorf/gorf.html). Comparisons of nucleotide and transcribed amino acid sequences were performed against public databases (NCBI) using BlastN, BlastX and BlastP (BLAST, basic local alignment search tool) at the NCBI on-line server. For translation to protein sequences, the bacterial code was selected, allowing ATG, GTG, and TTG as alternative start codons. All the predicted ORFs longer than 270 bp were translated and used as queries in BlastP. Sequences with significant matches were further analysed with psiBlast, and their putative function was annotated based on their similarities to sequences in the COG (Clusters of Orthologous Groups) and Pfam (Protein Families) databases. Predicted general cellular functions were annotated only for known ORFs based on the MultiFam classification (Serres et al, 2006). All sequences with an E-value higher than 0.001 in the BlastP searches and longer than 300 bp were considered to be unknown. Transmembrane helices were predicted with TMprep (http://www.ch.embnet.org/software/TMPRED_form.html) and signal peptides with Signal P3.0 server (http://www.cbs.dtu.dk/services/SignalP/). A complete table can be found at Supplementary Table S1 online. Promoter prediction was based on the analysis of the ten contigs by using both BPROM (http://www.softberry.com/berry.phtml?topic=bprom&group=programs&subgroup=gfindb) and bTSSfinder (http://www.cbrc.kaust.edu.sa/btssfinder/) web-based platforms. Both methods searched for rpoD-related sequences and we have only considered as valid predictions the ones matched on both approaches. Those filtered sequences were used to cross-validate 23 out of 33 experimentally defined regulatory regions by comparing the positions between predicted and experimental sequences in metagenomic fragments. The positions of the 33 small DNA fragments were obtained by a multiple alignment of the original contigs (queries) against those selected sequences, which has also allowed the validation of the promoter’s directionality – forward or reverse - by observing the matched strands (Plus/Plus or Plus/Minus). The consensus Logo sequence was based on the alignment of the 33 experimentally validated promoters, using the WebLogo platform (http://weblogo.berkeley.edu/logo.cgi).

### Calculations for promoter/Kb rates from databases and for promoter accessibility estimation

Data from predicted sequences of promoter sites, TSS (Transcriptional Start Site) and TUs (Transcription Unit) reported in different studies and databases regarding *E. coli* and other bacteria ^3,61,65,70^ were used as proxies for the total number of predicted promoters. Those values were divided by their respective genome sizes (or average genome sizes when calculating an average rate of multiple species at once) in order to provide promoter/Kb rates (i.e. 8,000 predicted promoters, TSS or TUs on a genome of 4.6 Mb would result in a rate of 1.7 promoters/Kb).The promoter accessibility estimation followed the same rationale and was based on the combination of the data from both metagenomics libraries presented in Table 1 and the rate of experimentally discovered promoters per Kb (33 promoters found in 30 Kb of metagenomic DNA, resulting in a rate of 1.1 promoters/Kb). Firstly, we have merged data from both metagenomic libraries and calculated the predicted number of promoters in a metagenomics library with an effective size of 503 Mb (combined effective sizes of USP1 and USP3) – the “effective size” takes into account only the percentage of clones with an insert -. Thus, we have multiplied the effective library size by the 23previously obtained promoter rate (1.1 promoters/Kb), resulting in a total estimated set of 553,300 promoter sequences. Secondly, we have calculated the predicted set of accessible promoters by multiplying the number of fluorescent clones (1,100 clones, considering both libraries) by the average insert size (4.1 Kb) and by the rate of observed promoters per Kb (1.1 promoters/Kb), resulting in 4,961 potentially accessible promoters. Lastly, we have calculated the proportion of accessible promoters among the total number of predicted promoters, which represents approximately ~1% of the whole available set.

### Data Availability

The nucleotide sequences obtained for the plasmid inserts have been deposited in the GenBank database under the Accession numbers (KY939589-KY939597), which are also shown in Table 2.

## Acknowledgements

This work was supported by the National Council for Technological and Scientific Development (CNPq 472893/2013-0 and 441833/2014-4) and by Young Research Awards by the Sao Paulo State Foundation (FAPESP, award numbers 2015/04309-1 and 2012/21922-8). CAW and LFA are beneficiaries of FAPESP fellowships (award numbers 2016/05472-6 and 2016/06323-4, respectively). Authors have no conflict of interest to declare.

## Author Contributions

CAW, LFA, MEG and RSR designed the experiments. CAW and LFA performed the experiments. CAW analyzed the data. CAW and RSR prepared the figures. CAW and MEG wrote the manuscript. All authors reviewed the manuscript.

